# Thiol-linked alkylation for the metabolic sequencing of RNA

**DOI:** 10.1101/177642

**Authors:** Veronika A. Herzog, Brian Reichholf, Tobias Neumann, Philipp Rescheneder, Pooja Bhat, Thomas R. Burkard, Wiebke Wlotzka, Arndt von Haeseler, Johannes Zuber, Stefan L. Ameres

## Abstract

Gene expression profiling by high-throughput sequencing reveals qualitative and quantitative changes in RNA species at steady-state but obscures the intracellular dynamics of RNA transcription, processing and decay. We developed thiol(SH)-linked alkylation for the metabolic sequencing of RNA (SLAM-seq), an orthogonal chemistry-based epitranscriptomics-sequencing technology that uncovers 4-thiouridine (s^4^U)-incorporation in RNA species at single-nucleotide resolution. In combination with well-established metabolic RNA labeling protocols and coupled to standard, low-input, high-throughput RNA sequencing methods, SLAM-seq enables rapid access to RNA polymerase II-dependent gene expression dynamics in the context of total RNA. When applied to mouse embryonic stem cells, SLAM-seq provides global and transcript-specific insights into pluripotency-associated gene expression. We validated the method by showing that the RNA-polymerase II-dependent transcriptional output scales with Oct4/Sox2/Nanog-defined enhancer activity; and we provide quantitative and mechanistic evidence for transcript-specific RNA turnover mediated by post-transcriptional gene regulatory pathways initiated by microRNAs and *N*^6^-methyladenosine. SLAM-seq facilitates the dissection of fundamental mechanisms that control gene expression in an accessible, cost-effective, and scalable manner.

**One Sentence Summary:** Chemical nucleotide-analog derivatization provides global insights into transcriptional and post-transcriptional gene regulation

## Introduction

The regulated expression of genetic information imperatively stipulates cellular homeostasis and environmental adaptability and its transformation entails human diseases, such as cancer^1^. Underlying these fundamental biological processes are tightly regulated molecular events that control the relative kinetics of RNA transcription, processing, and degradation. Understanding the molecular basis for gene regulatory circuits demands detailed insights into the relative kinetics of RNAs biogenesis and degradation in a transcript-specific and systematic manner^2^.

With the advent of high-throughput sequencing technologies, several approaches have been developed to globally address the kinetics of gene expression at the genomic scale. For example, genome-wide analyses of nascent RNAs engaged with transcriptionally active RNA polymerase by high throughput sequencing (e.g. GRO-seq, PRO-seq, NET-seq) represent powerful methods to investigate transcriptional activity^3-5^. Despite their utility, these approaches are technically challenging and do not discriminate functional and processed RNA molecules from transcriptional by-products. On the other hand, RNA decay rates are commonly determined by globally interfering with RNA polymerase II (Pol II) activity followed by the relative quantification of RNA molecules over time^6^. But due to the global loss of RNAs, the quantification of absolute RNA half-lives remains imprecise. Furthermore, inhibiting polymerase II activity compromises cell viability and induces major stress responses, triggering secondary effects including the selective stabilization of specific transcripts^7-10^.

Metabolic RNA labeling approaches that employ the incorporation of nucleotide analogs enable tracking of RNA species over time without interfering with cellular integrity. Among these, 4-thiouridine (s^4^U) represents the most widely used nucleotide analog to study the dynamics of RNA expression because it is readily imported into metazoan cells by equilibrate nucleoside transporters^11^, and provides unique physicochemical properties for thiol-specific reactivity and affinity, which enables the biochemical separation by reversible biotinylation [e.g. through N-[6-(Biotinamido)hexyl]-3′-(2′-pyridyldithio)propionamide (HPDP-Biotin) or biotin-coupled methanethiosulfonates (MTS-Biotin)]^12-16^. Affinity-based RNA-purification upon s^4^U-labeling has been successfully applied to cultured cells of diverse biological and organismal origin, as well as in vivo in yeast and metazoan model organisms, including insects and mammals, using either 4-thiouridine or 4-thiouracil upon metabolic activation by uracil phosphoribosyltransferase (UPRT)^12,13,17-19^. However, like any biochemical separation method, the underlying protocols are laborious, require ample starting material, and typically encounter the problem of low signal-to-noise performance, in part because of limited biotinylation efficiency^15^. Furthermore, the analysis of labeled RNA species by sequencing requires extensive controls in order to provide integrative insights into gene expression dynamics and fails to report global effects unless spike-in strategies are applied ^16,20^. Alternative concepts for the direct identification of nucleotide analogs by sequencing emerge from recent epitranscriptomics-technologies that uncover RNA modifications by orthogonal chemistry and sequencing, but current methods are incompatible with biologically inert nucleotide-analogs (i.e. s^4^U) and fail to report absolute stoichiometry^21,22^.

Here, we report thiol(SH)-linked alkylation for the metabolic sequencing of RNA (SLAM-seq), an orthogonal chemistry approach that uncovers s^4^U-incorporation events at single-nucleotide resolution by reverse-transcription-dependent thymine-to-cytosine-conversions in a high-throughput sequencing-compatible manner. By combining thiol-linked alkylation with s^4^U-metabolic RNA labeling in mouse embryonic stem cells followed by mRNA 3′ end sequencing, SLAM-seq provides global and transcript-specific insights into RNA polymerase II-dependent, pluripotency-associated gene expression and its regulation at the transcriptional and post-transcriptional level. SLAM-seq provides a powerful tool for the dissection of fundamental biological mechanisms that control the implementation of genetic information.

## Results

### Detection of 4-thiouridine by sequencing

To provide rapid and direct experimental access to metabolically labeled RNA by sequencing, we developed a chemical-derivatization approach for 4-thiouridine (s^4^U), a nucleotide-analog widely used for in vivo and ex vivo RNA labeling (Fig.1). To this end, we employed the primary thiol-reactive compound iodoacetamide (IAA), which covalently attaches a carboxyamidomethyl-group to s^4^U by nucleophilic substitution (Fig.1a). Quantitative s^4^U-alkylation was confirmed by a shift in the characteristic absorbance spectrum of 4-thiouracil from ∼335 nm to ∼297 nm (Fig.1b)^23^. Under optimal reaction conditions (Supplementary Fig.1), absorbance at 335 nm decreased 50-fold compared to untreated 4-thiouracil, resulting in complete (≥98%) alkylation within 15 min (Fig.1c and Supplementary Fig.1). Mass spectrometry analysis of thiol-specific alkylation in a ribose-context confirmed these derivatization-efficiencies (Fig.1d, e, and Supplementary Fig.2). Because quantitative identification of s^4^U by sequencing presumes that reverse transcriptase (RT) passes alkylated s^4^U-residues without drop-off, we determined the effect of s^4^U-alkylation on RT-processivity in primer extension assays (Fig.1f). We did not observe a significant effect of s^4^U-alkylation on RT processivity when compared to a non-s^4^U-containing oligo with identical sequence (Fig.1g and Supplementary Fig.3). To evaluate the effect of s^4^U-alkylation on RT-directed nucleotide incorporation, we isolated the full-length products of primer extension reactions, PCR-amplified the cDNA and subjected the libraries to high-throughput sequencing (Fig.1h, i, and Supplementary Fig.4). While the presence of s^4^U prompted a constant ten to eleven percent T>C conversions already in the absence of alkylation (presumably due to base-pairing variations of s^4^U-tautomeres), s^4^U-alkylation increased T>C conversions by 8.5-fold, resulting in a >0.94 conversion rate (Fig.1i). Importantly, iodoacetamide-treatment leaves conversion rates of any given non-thiol-containing nucleotide unaltered (Supplementary Fig.4c). We concluded that iodoacetamide treatment followed by reverse transcription specifically identifies s^4^U-incorporations in RNA at single-nucleotide resolution by sequencing at >90% efficiency.

**Figure 1.**
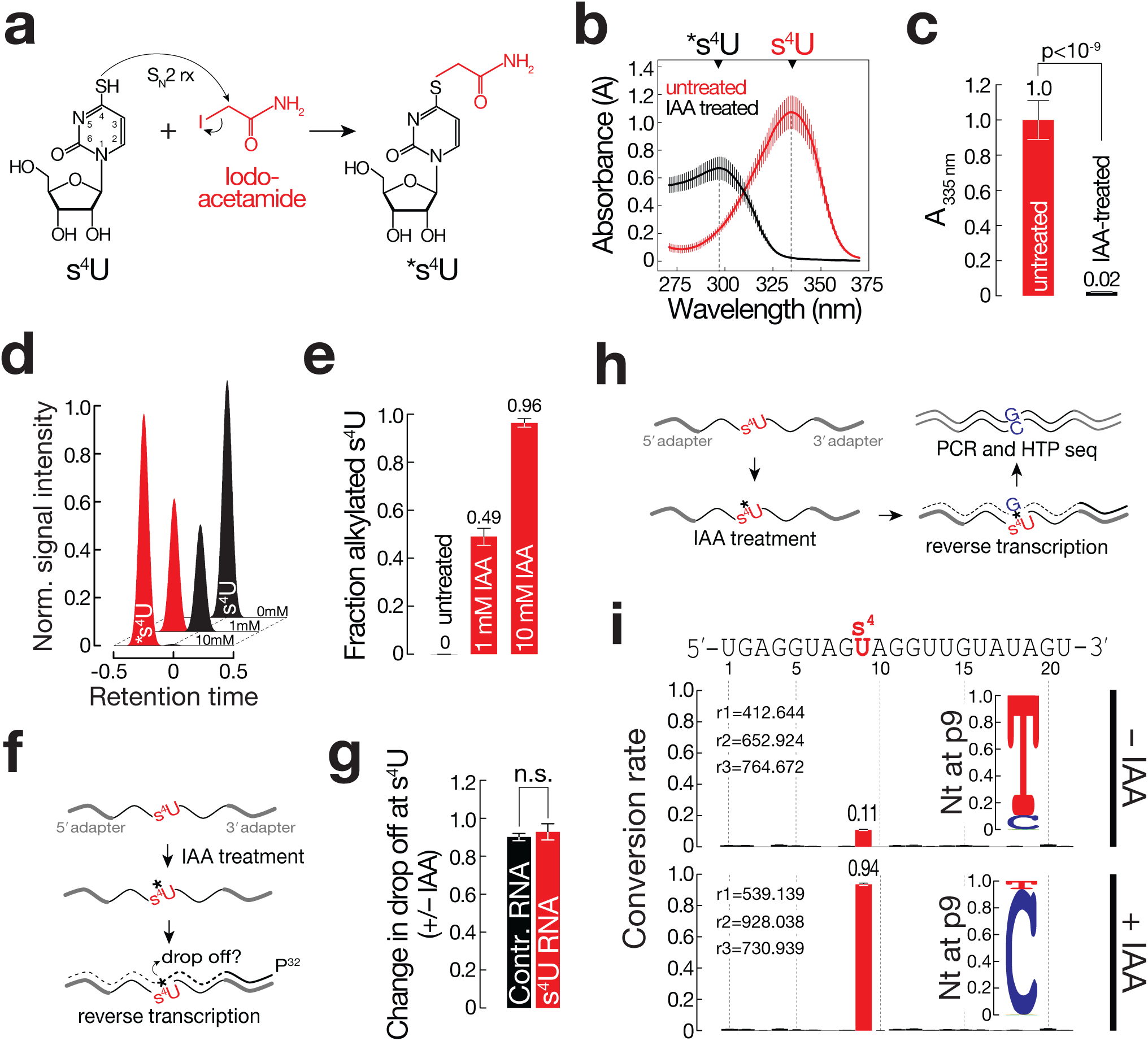
Detection of 4-thiouridine (s^4^U) by chemical derivatization and sequencing. **(a)** 4-thiouridine (s^4^U) reacts with the thiol-reactive compound iodoacetamide (IAA), attaching a carboxyamidomethyl-group to the thiol-group in s^4^U as a result of a nucleophilic substitution (S_N_2) reaction. **(b)** Absorption spectra of 4-thiouracil (s^4^U) before and after treatment with iodoacetamide (IAA). Absorption maxima of educt (4-thiouracil; s^4^U; λ_max_ ≈ 335 nm) and product (carboxyamido-methylated 4-thiouracil; *s^4^U; λ_max_ ≈ 297 nm) are indicated. Data represents mean ± SD of at least three independent replicates. Quantification of absorption at 335 nm as shown in (b). Data represents mean ± SD of at least three independent replicates. P-value (two-tailed Student’s t-test) is indicated. Normalized LC-MS extracted ion chromatograms of s^4^U (black) and alkylated s^4^U (red) at the indicated iodoacetamide concentrations. **(e)** Quantification of two independent experiments shown in (d). Fraction alkylated s^4^U at indicated IAA concentrations represent relative normalized signal intensities at peak retention times of s^4^U and alkylated s^4^U. Data represent mean ± SD of two independent experiments measured in two technical replicates. **(f)** The effect of s^4^U-alkylation on reverse transcriptase-processivity was determined by primer extension assay. **(g)** Quantification of three independent replicates of experiment shown in (f). Ratio of drop off signal (+ vs – IAA treatment) at s^4^U-position after normalization to preceding background drop off signal was determined for control and s^4^U-containing RNA. Data represent mean ± SD. P-value (two-tailed Student’s t-test) is indicated. **(h)** RNA with or without 4-thiouridine (s^4^U) incorporation at a single position (p9) was treated with iodoacetamide (IAA) and subjected to reverse transcription and gel-extraction of full-length product followed by PCR amplification and high-throughput (HTP) sequencing. **(i)** Conversion rates for each position of a s^4^U-containing RNA before or after iodoacetamide (IAA) treatment. Average conversion rates ± SD of three independent replicates are shown. Number of sequenced reads in each replicate (r1-r3) are indicated. Nucleotide identity at s^4^U site (p9) is shown.

### SLAM-seq quantifies s^4^U-labeled transcripts in mES cells

To employ thiol-linked alkylation for the metabolic sequencing of RNA (SLAM-seq; Fig.2a), we subjected mouse embryonic stem (mES) cells to s^4^U-labeling at 100 μM, a concentration far below the EC_50_ toxicity value of s^4^U in mES cells (882 μM after repeated exposure of mES cells to s^4^U in 3h intervals for 24 h; Supplementary Fig.5). After metabolic RNA labeling for 24 h, we prepared total RNA followed by thiol-alkylation and 3′ end mRNA sequencing (Quant-seq). Quant-seq provides rapid and quantitative access to mRNA expression profiles from low quantities of total RNA (0.5-500 ng)^24^, by generating Illumina-compatible libraries of the sequences close to the 3′ end of polyadenylated RNA (Fig.2b, Supplementary Fig.6). In contrast to other mRNA-sequencing protocols, only one fragment per transcript is generated, which corresponds to polyadenylated mRNA 3′ end tags (Supplementary Fig.6), rendering normalization of reads to gene length obsolete. This results in accurate, highly-reproducible, and strand-specific gene expression values (Supplementary Fig.6)^24^. Furthermore, 3′ end sequencing enables the cell-type-specific re-evaluation of UTR-annotations to conduct mRNA 3′ isoform-specific expression analysis (Supplementary Fig.7). Upon generating SLAM-seq libraries through the Quant-seq protocol from total RNA of mES cells 24h after s^4^U metabolic labeling, we observed a strong accumulation of T>C conversions when compared to libraries prepared from total RNA of unlabeled mES cells (Fig.2b). Transcriptome-wide alignment of reads to mRNA 3′ ends confirmed this observation (Fig.2c): In the absence of s^4^U metabolic labeling, we observed a median rate of ≤0.1% for any given conversion, consistent with Illumina-reported sequencing error. Metabolic labeling with s^4^U resulted in a statistically significant (p<10^-4^, Mann-Whitney test), >50-fold increase in T>C conversion rates (Fig.2c), which distributed evenly across the covered genomic regions (Fig.2d, and Supplementary Fig.8). In contrast, non-T>C conversions remained below the expected sequencing error rates (Fig.2c). Importantly, treatment of total RNA with iodoacetamide in the absence of metabolic labeling did not affect quantitative gene expression analysis (Supplementary Fig.8d).

**Figure 2.**
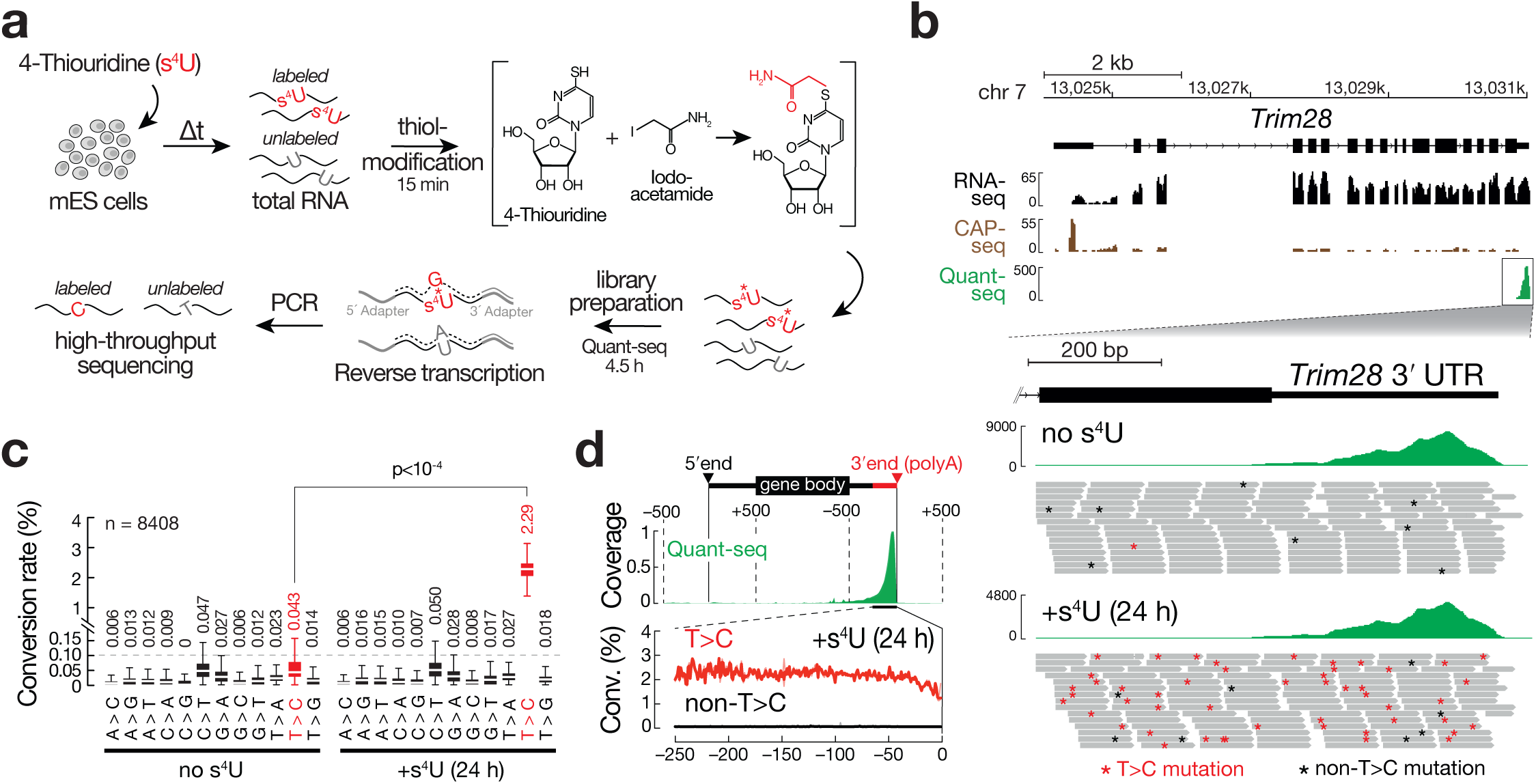
Thiol-linked alkylation for the metabolic sequencing of RNA (SLAM-seq). **(a)** Workflow of SLAM-seq. Working time for alkylation and Quant-seq library preparation are indicated. **(b)** Representative genome browser screen shot for three independent mRNA libraries generated from total RNA of mES cells, prepared using standard mRNA sequencing (top panel), Cap-seq (middle panel) and mRNA 3′ end sequencing (bottom panel; RPM, reads per million,). A representative area in the mouse genome encoding the gene Trim 28 is shown. Bottom shows zoom into 3′ UTR of Trim28. Unnormalized coverage plots of Quant-seq libraries prepared from untreated mES cells or mES cells subjected to s^4^U-metabolic labeling using 100 μM s^4^U for 24 h followed by SLAM-seq. A subset of individual reads underlying the coverage plots are depicted. Asterisks indicate T>C conversions (red) or any conversion other than T>C (black). **(c)** Conversion rates in 3′ UTR-mapping reads of Quant-seq libraries, prepared from mES cells before (no s^4^U) and after metabolic labeling for 24 h using 100 μM s^4^U (+s^4^U). Dashed line represents expected background sequencing error rate. Median conversion rate across the indicated number of transcripts (n) is indicated. P-value (Mann-Whitney test) is indicated. **(d)** Relative coverage across transcripts for mRNA-seq and Quant-seq. T>C conversion rate (Conv.) distributes evenly within Quant-seq-covered areas.

Together with the fact that s^4^U incorporation measured by mass spectrometry in poly(A)-enriched RNA and SLAM-seq were comparable even after shorter labeling times (i.e. 6h, Supplementary Fig.9), these analyses confirm that SLAM-seq provides robust access to s^4^U-incorporation events in mRNA following s^4^U-metabolic RNA labeling in cultured cells.

### Measuring the polyadenylated transcriptional output in mES cells

Steady-state gene expression is dictated by the relative contributions of the rate of transcription and RNA stability, two parameters that cannot be derived from standard RNA sequencing approaches. To test if SLAM-seq provides quantitative access to the polyadenylated transcriptional output, we subjected mES cells to 45 min s^4^U-pulse labeling (final conc.:100 μM s^4^U) followed by total RNA extraction, alkylation, and mRNA 3′ end library preparation (Fig.3a). We approximated the number of molecules that were made within the short timeframe of labeling by inspecting background-error-subtracted T>C conversion-containing reads for individual transcripts. Indeed, initial inspection of selected transcripts with comparable steady-state abundance (∼100 cpm) revealed transcript-specific differences in the number of recovered T>C reads (Fig.3b): While high levels of T>C reads were recovered for the ES cell-specific transcription factor Sox2, and the inherently instable primary microRNA transcript from the miR-290-295 cluster, the house-keeping transcript Gapdh associated with fewer T>C reads, presumably because its accumulation to high steady-state expression levels is achieved by high transcript stability (Fig.3b). (A global overview of transcriptional output measurements by SLAM-seq is provided in Supplementary Table 1.)

**Figure 3.**
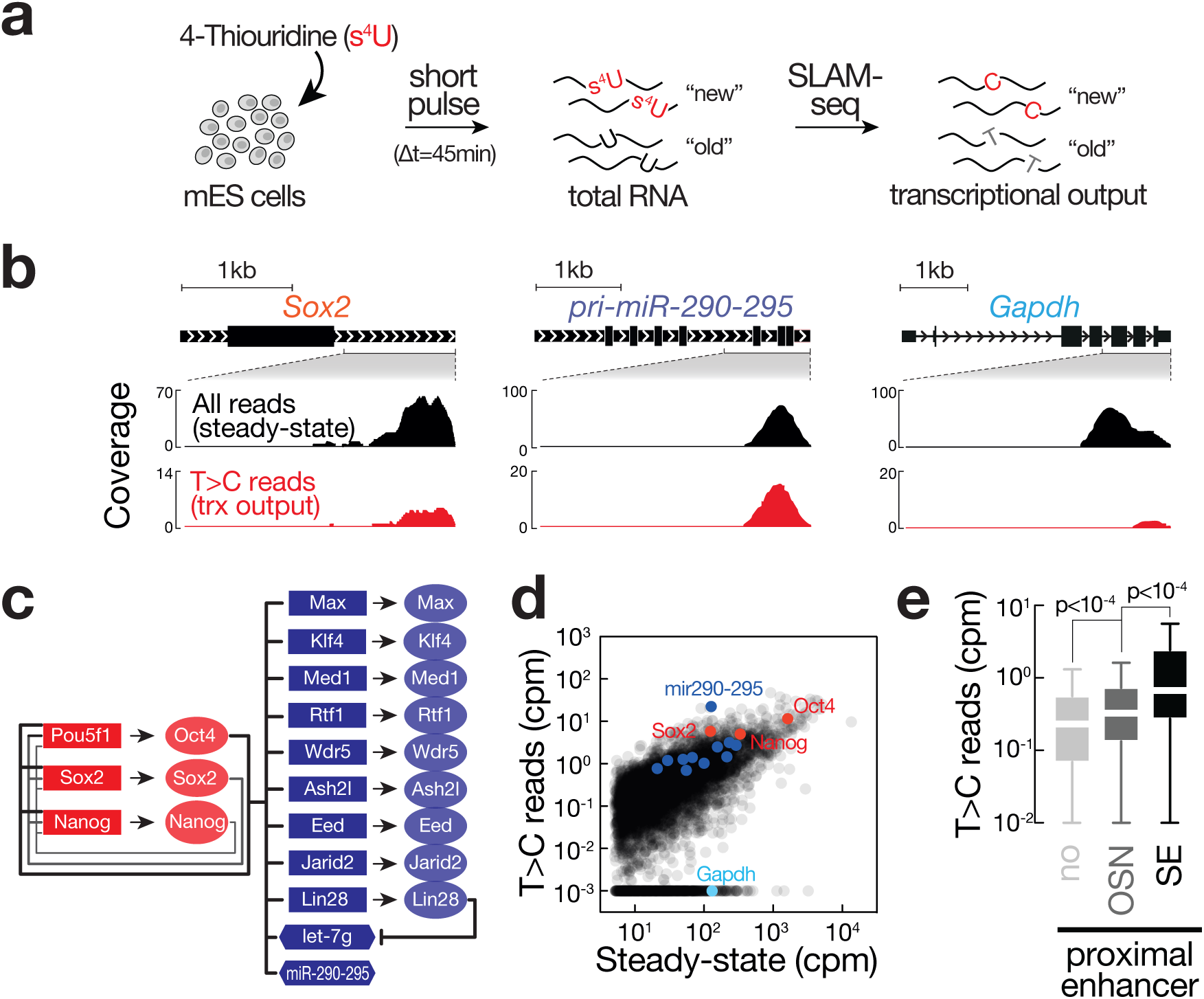
Quantitative description of the polyadenylated transcriptional output in mES cells. **(a)** Experimental setup for determining transcriptional mRNA output by SLAM-seq, coupled to short (45 min) s^4^U-pulse labeling using 100 μM s^4^U. **(b)** Genome browser plots of the indicated genes show SLAM-seq data prepared from mES cells, subjected to s^4^U-metabolic RNA labeling as shown in (a). Black reads represent all mapped reads (steady-state, in RPM); red reads represent T>C conversion-containing reads (de novo transcribed; trx output, in RPM). **(c)** Model of genes involved in maintenance of stem cell state (adapted from^26^). **(d)** Relative transcriptional output for 7179 genes in mES cells. T>C reads represent abundance of de novo transcripts in counts per million (cpm); Steady-state represents sum of T>C- and non-T>C-containing reads. Core pluripotency transcription factors are highlighted in red, a subset of primary target genes (see c) in blue and a gene with house-keeping function in light-blue. **(e)** Transcriptional output, as measured in number of T>C conversion containing reads, for expressed genes (steady-state >5cpm) without adjacent OSN enhancer (no, n=4994), proximal to canonical Oct4/Sox2/Nanog enhancer (OSN, n=2029) or proximal to strong enhancers (SE, n=156). Outliers are not shown. P-values determined by Mann-Whitney test are indicated.

Transcriptional output by Pol II is regulated by transcription factors that bind cis-acting regulatory elements known as enhancers to recruit coactivators and Pol II to target genes^25^. In ESCs the pluripotent state is largely governed by a small number of master transcription factors, including Oct4, Sox2, and Nanog, which drive the expression of target genes necessary to maintain the ESC state and positively regulate their own promoters, forming an interconnected auto-regulatory loop (Fig.3c)^26^. Transcriptional output measurements by SLAM-seq revealed that Oct4/Sox2/Nanog target genes produced overall a larger number of T>C conversion containing reads in the s^4^U-pulse experiment, confirming their high transcriptional activity (Fig.3d, Supplementary Table 1). To connect these observations to molecular principles of transcriptional gene regulation, we correlated the measured transcriptional output to proximal enhancer loci (Fig.3e), which were classified by their association with the transcription factors Oct4/Sox2/Nanog (OSN enhancer; OSN), as previously described^27^. Transcripts derived from the 2029 expressed genes (>5cpm steady-state) with proximal OSN occupancy produced significantly higher levels of T>C reads upon pulse labeling when compared to 4994 genes without proximal OSN enhancer (Mann-Whitney test, p<10^-4^). A subset of enhancers in mES cells were previously described to form arrays of concatenated regulatory elements (aka “super” or strong enhancer, SE), with unusually strong accumulation of transcriptional coactivators, specifically Mediator^27,28^. In fact, the 156 genes in the proximity of such strong enhancers exhibited highest transcriptional output, significantly off-set from typical enhancers in our SLAM-seq measurements (Mann-Whitney test, p<10^-4^, Fig 3e). In contrast, only the genes proximal to strong enhancers associated with above average expression at steady-state (Supplementary Fig.10). We concluded that SLAM-seq provides a quantitative readout for enhancer activity-associated transcription in mES cells.

Together with the fact that the obtained transcriptional output measurements significantly correlated with data derived from global nuclear run-on experiments (Spearman’s correlation r_s_=0.41, p<10^-15^, Supplementary Fig.11)^29^, we concluded that short s^4^U-pulse labeling in combination with SLAM-seq and mRNA 3′ end sequencing enables to uncouple the immediate transcriptional output from stability effects to directly measure the functional Pol II-derived transcriptional activity at the genomic scale. The method therefore provides a rapid and scalable approach to study transcriptional gene regulatory circuits in mammalian cells.

### Global and transcript-specific mRNA stability in mES cells

To apply SLAM-seq for direct measurements of mRNA transcript stabilities we subjected mES cells to s^4^U metabolic RNA labeling (100 μM s^4^U) for 24 h, followed by washout and chase using non-thiol-containing uridine, and prepared total RNA at various time points along the chase (0h, 0.5h, 1h, 3h, 6h, 12h, and 24h). Total RNA was then subjected to alkylation and mRNA 3′ end sequencing (Fig.4a). Inspection of candidate genes revealed constant steady-state expression across the whole time-course (Fig.4b, all reads). In contrast, T>C-conversion containing reads decreased over time in a transcript-specific manner (Fig.4b, T>C reads). After calculating the background-subtracted, U-content- and coverage-normalized T>C conversion rate for each transcript at every time point relative to 0h chase, normalized T>C conversion rates fit well to single-exponential decay kinetics, enabling the robust determination of polyadenylated transcript half-lives (t_½_). As expected, RNA stabilities differed by more than one order of magnitude among individual transcripts (Fig.4c). A short half-life was observed for the primary miRNA transcript of the miR-290-295 cluster (t_½_=0.8h), most certainly because it is subjected to rapid co-transcriptional processing by the RNase III enzyme Drosha in the nucleus^30^. To validate these results globally, we first determined the relative steady-state gene expression profiles along the chase experiment and detected high overall correlation (Spearman’s correlation r_S_>0.9), while correlations between the abundance of T>C conversion containing transcripts and steady-state decreased over time (Fig.4d top). This was expected, since transcript-specific stabilities are not dependent on steady-state abundance. By fitting the data of 8405 transcripts with a steady-state expression cutoff of >5 reads per million to single-exponential decay kinetics, we determined a median mRNA half-life of 3.9h, corresponding to a cell-cycle normalized half-life of 4.3h (Fig.4d left, Supplementary Table 2). These measurements fall within the range of previously determined mRNA stabilities in mammalian cells, ranging in median half-life from 3.4h to 10h, depending on cell type and experimental technique^6^.

**Figure 4.**
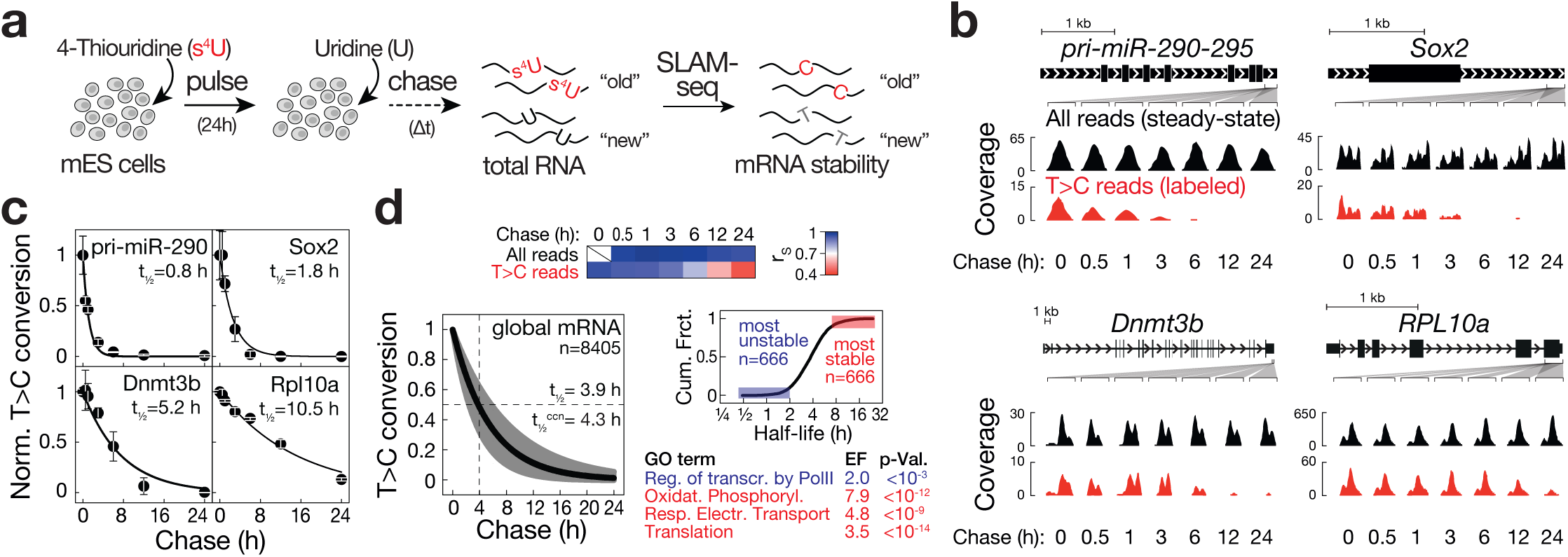
Global and transcript-specific mRNA stability in mES cells. **(a)** Experimental setup for profiling mRNA stability by SLAM-seq. mES cells were subjected to metabolic labeling with s^4^U (100 μM final conc.) in 3h intervals for a total of 24 h, followed by a chase with unlabeled uridine (10 mM final conc.) for 0, 0.5, 1, 3, 6, 12, and 24 h, followed by total RNA preparation and SLAM-seq. **(b)** Genome browser plots of the indicated genes represent SLAM-seq data prepared from mES cells subjected to s^4^U-pulse/chase labeling as shown in (a). All mapped reads (steady-state, in RPM) are black; T>C conversion-containing reads (labeled, in RPM) are red. **(c)** Transcript stability for the indicated genes. T>C-conversion rates were determined for each timepoint of the s^4^U-pulse/chase and fit to a single-exponential decay model to derive half-life (t_½_). Values are mean ± SD of three independent experiments. **(d)** Global analysis of mRNA stability in mES cells. Top: Correlation of steady-state gene expression (all reads) or T>C conversion containing reads at the indicated time with steady-state expression at time 0. Left bottom: RNA half-life for 8405 transcripts in mES cells determined as described in (c). Median half-life before (t_½_) or after (t^ccn^_½_) normalization to cell divisions. Right bottom: Cumulative distribution of ranked transcript stabilities for 6665 transcripts. Enriched gene ontology (GO) terms for the 666 most unstable (blue) or most stable (red) are indicated. Enrichment factor (EF) and p-value (p-Val.) are indicated.

Previous genome-wide investigations of mRNA stability proposed a close relationship between transcript-specific mRNA half-life and its physiological function^1,18^. We therefore ranked the 6665 transcripts for which half-life was determined at high accuracy (r^2^>0.6) according to their relative stability and performed gene ontology enrichment analysis for the 666 most or least stable mRNAs (Fig.4d right panel). Transcripts with short half-life significantly enriched for regulators of Pol II-dependent transcription (p<10^-3^), while stable mRNAs associated with the GO terms translation (p<10^-14^), respiratory electron transport (p<10^-9^) and oxidative phosphorylation (p<10^-12^). Together with gene set enrichment analyses (Supplementary Fig.12), SLAM-seq measurements confirmed that transcripts encoding proteins with house-keeping function, such as protein synthesis or respiration, tend to decay at low rates, perhaps reflecting the evolutionary adaptation to energy constraints. In contrast, transcripts with a regulatory role, such as transcription factors or cell-cycle genes, tend to decay faster, most certainly because control over the persistence of genetic information facilitates adaptation to environmental changes^1^.

We also examined global relationships between transcriptional output, mRNA stability, and steady-state gene expression in mES cells as determined by SLAM-seq pulse and pulse/chase experiments (Supplementary Fig.13): We found that transcript biogenesis rates and mRNA half-life both positively correlated with steady-state gene expression with correlation coefficients of 0.57 and 0.43, respectively. In contrast, the rates of mRNA biogenesis did not positively correlate with mRNA half-life (r=-0.07), but showed high correlation with mRNA decay rates (r=0.66). These results agree with a transcript-specific contribution of both mRNA synthesis and decay to the establishment of steady-state gene expression in mES cells, hence validates the applicability of SLAM-seq to studying both transcriptional and post-transcriptional gene regulatory mechanisms.

Together with the fact that mRNA half-life measurements showed an overall good correlation with mRNA stabilities determined in mES cells after transcription-inhibition using Actinomycin D (r=0.77, Supplementary Fig.14), we concluded that s^4^U-pulse/chase labeling in combination with SLAM-seq and mRNA 3′ end sequencing provides rapid access to global and transcript-specific mRNA stability.

### SLAM-seq uncovers molecular determinants of mRNA stability

To further validate SLAM-seq, we performed mechanistic studies on two specific post-transcriptional gene regulatory pathways with well-established biological functions in mES cells that have previously been implicated in transcript-specific mRNA decay:

First, we focused on microRNAs (miRNAs), which act as key regulators of gene expression in a variety of biological contexts^31^. In mES cells, they contribute to cell state maintenance and transitions by tuning the expression of ES cell transcripts and promoting their clearance during differentiation^26^. At the molecular level, miRNAs act as guides for ribonucleoprotein complexes that target complementary sites, usually within the 3′ UTR of mRNAs^31^. Target-binding is primarily mediated via the miRNA seed sequence, encompassing nucleotides two to seven or eight when counted from the miRNA 5′ end. MicroRNAs elicit their function by repressing translation and/or promoting mRNA decay, although the relative contribution of repressive modes remains a matter of debate and may vary in different biological contexts (Fig.5a) ^32^. To estimate the effect of miRNAs on mRNA stability in mES cells, we first determined the general association of miRNA targets with mRNA stability in wild-type cells by inspecting half-life of transcripts harboring in their 3′ UTR target sites for the miR-291-3p/294-3p/295-3p/302-3p and miR-292a-3p/467a-5p-family (in the following referred to as miR-291a-family) (Fig.5b). Members of the miR-291a-family share the same seed sequence and target-repertoire, and derive from the ESC-specific miR-290-295 cluster that gives rise to more than half of all small RNAs expressed in this cell type (Supplementary Fig.15). With a median half-life of 2.9h (n=1450), miR-291a-family targets were significantly less stable compared to transcripts that contained no target sites (t_½_=4.0h; n=5095; KS-test, p<10^-15^; Fig.5b). Transcripts with conserved sites exhibited even shorter half-life (t_½_=2.6h; n=50; Fig.5b). To confirm the direct contribution of miRNAs to transcript destabilization, we determined changes in mRNA half-life by s^4^U-pulse labeling followed by SLAM-seq in mES cells depleted of the core miRNA biogenesis factor Exportin-5 (Xpo5) by CRISPR/Cas9 genome engineering (Fig.5c, Supplementary Fig.16a, b). Compared to wild-type cells, depletion of Xpo5 significantly decreased overall miRNA levels by more than 90%, and miR-291a-family members by more than 95%, as determined by small RNA sequencing (Student’s t-test p<10^-4^, Fig.5d) and confirmed by Northern hybridization (Supplementary Fig.16c). Consistent with a function of miRNAs in triggering mRNA decay, we observed a significant increase in relative mRNA stability for targets of the miR-291a-family when compared to transcripts containing no target site (KS-test, p<10^-15^ and p<10^-4^ for all or conserved sites, respectively; Fig.5e). Notably, the degree of de-repression followed previously established rules for miRNA targeting (Fig.5a and f)^31^: While each site-type responded to Xpo5-depletion with a significant increase in mRNA stability relative to untargeted transcripts (KS-test, p<10^-8^), 6mer target sites exhibited the weakest effects, followed the two 7mer site types (Fig.5f). 8mer sites showed strongest de-repression (Fig.5f). Finally, by inspecting target mRNAs of less abundant miRNA families we confirmed that miRNA function, as determined by target mRNA stabilities in wild-type mES cells and relief of repression upon depletion of Xpo5, is directly dependent on small RNA abundance (Supplementary Fig.17), as described previously ^33,34^. In summary, these data provide quantitative evidence for miRNA-mediated mRNA decay in mES cells and validate SLAM-seq as a sensitive tool to study the molecular basis underlying the fine-tuning of mRNA expression levels by miRNAs.

**Figure 5.**
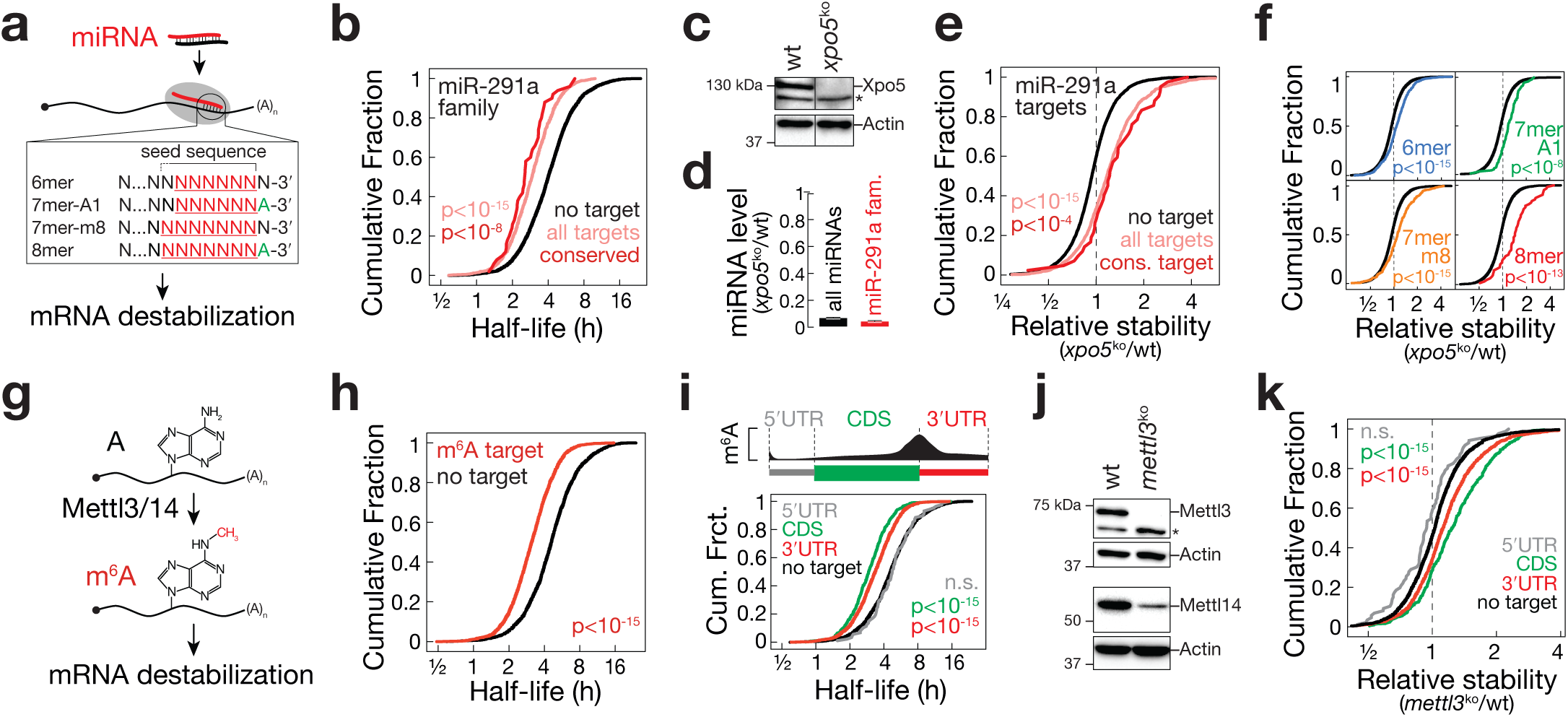
Molecular determinants of mRNA stability in mES cells. **(a)** Schematic representation of microRNA (miRNA)-mediated gene regulation. Target site-types with increasing (top to bottom) repressive function are shown. **(b)** Cumulative distribution of ranked mRNA stabilities. Plotted are distributions for transcripts that do (rose, n=1450) or do not (black, n=5095) contain at least one miR-291a-family target site or contain at least one conserved miR-291a target site (red, n=50). miR-291a-family members as defined in Supplementary Fig.17a. P-value was determined by KS-test. **(c)** Western blot analysis for Exportin-5 (Xpo5) and Actin in wild-type (wt) mES cells and a cell line depleted of Xpo5 by CRISPR/Cas9-induced frameshift in the first coding exon (see Supplementary Fig.16). **(d)** MicroRNA ratio determined by spike-in-controlled small RNA sequencing in *xpo5*^ko^ mES cells relative to wild-type cells. Ratio for all miRNAs (black) or miR-291a-family members (red) are shown. **(e)** Cumulative distribution of mRNA stability changes in *xpo5*^ko^ relative to wt mES cells. Plotted are distributions for transcripts that do (rose, n=1450) or do not (black, n=5095) contain at least one miR-291a target site or that contain at least one conserved miR-291a target site (red, n=50). P-value was determined by KS-test. **(f)** Cumulative distribution of mRNA stability changes in *xpo5*^ko^ relative to wt mES cells. Plotted are distributions for transcripts that contain exclusively one 6mer (blue, n=493), 7mer-A1 (green, n=95), 7mer-m8 (yellow, n=325), or 8mer site (red, n=63). Black shows transcripts without any miR-291a target site (n=5095). P-value was determined by KS-test. **(g)** Schematic representation of adenosine (A) conversion into *N6*-methyladenosine (m^6^A) by the methyltransferase complex Mettl3/14. **(h)** Cumulative distribution of ranked mRNA stabilities. Plotted are distributions for transcripts that do (red, n=3492) or do not (black, n=3173) contain the m^6^A mark, as previously mapped by m^6^A-RIP-seq^37^. P-value was determined by KS-test. **(i)** Top: Schematic distribution of m^6^A within mRNA (adapted from^40^). Bottom: Cumulative distribution of ranked mRNA stabilities. Plotted are distributions for transcripts that do not (black, n=3173) or do contain m^6^A exclusively in the 5′ UTR (grey, n=88), the coding sequence (CDS, green, n= 545) or the 3′ UTR (red, n=2093). P-value was determined by KS-test. **(j)** Western blot analysis for Mettl3, Mettl14 and Actin in wild-type (wt) mES cells and a cell line depleted of Mettl3 by CRISPR/Cas9-induced frameshift in the first coding exon (see Supplementary Fig.16). **(k)** Cumulative distribution of mRNA stability changes in *mettl3*^ko^ relative to wt mES cells. Plotted are distributions for transcripts that do not (black, n=3173) or do m^6^A exclusively in the in the 5′ UTR (grey, n=88), the coding sequence (CDS, green, n= 545) or the 3′ UTR (red, n=2093). P-value was determined by KS-test.

Second, we focused on *N*^*6*^-methyladenosine (m^6^A), the most abundant internal modification in mammalian mRNA, implicated in the regulation of various physiological processes^35,36^. In ESCs, m^6^A facilitates the resolution of naïve pluripotency towards differentiation^37,38^. At the mechanistic level, the m^6^A mark impinges on various aspects of mRNA processing, including mRNA stability^39^ (Fig.5g). To estimate the effect of m^6^A on mRNA stability in mES cells, we first determined the general association of m^6^A targets, as mapped previously by m^6^A-RNA-immunoprecipitation and sequencing^37^, with mRNA stability in wild-type cells. With a half-life of 3.1h, m^6^A-containing transcripts (n=3492) were significantly less stable compared to naïve transcripts (t_½_=4.6h, n=3173, KS-test, p<10^-15^, Fig.5h). *N*^*6*^-methyladenosine marks do not distribute evenly within mRNAs but are enriched in long exons, near stop codons, and in 3′ untranslated regions (UTRs), although m^6^A also occurs in the coding region (CDS) and 5′ UTR^40^ (Fig.5i top). We therefore investigated the relationship between the position of m^6^A within targeted mRNAs and its impact on RNA decay. We found that mRNAs containing m^6^A exclusively in the CDS (n=545) or in the 3′ UTR (n=2093) were significantly less stable compared to naïve transcripts (KS-test, p<10^-15^, Fig.5i bottom). In contrast, mRNAs that contained m^6^A exclusively in the 5′ UTR (n=88) were not less stable compared to naïve transcripts (KS-test, p>0.05, Fig.5i). To confirm the causal contribution of m^6^A to transcript destabilization we determined changes in mRNA half-life by s^4^U-pulse labeling followed by SLAM-seq in mES cells depleted of Mettl3, the catalytic subunit of the m^6^A RNA methylation complex, which resulted in the co-depletion of its RNA-binding partner protein Mettl14 (Fig.5j and Supplementary Fig.16d, e)^39^. Consistent with a direct and position dependent impact of m^6^A on mRNA decay, we observed a significant increase in relative mRNA stability for transcripts containing m^6^A in the CDS or 3′ UTR (KS-test, p<10^-15^) but not in the 5′ UTR (p>0.05; Fig.5k). Similar results were obtained when re-investigating recently described m^6^A profiling data in mES cells (Supplementary Fig.18)^41^. Taken together, our results provide quantitative evidence for m^6^A-mediated mRNA decay in mES cells and validate SLAM-seq as a sensitive method to study the function of chemical RNA modifications in RNA decay.

### Discussion

Recent efforts in decoding RNA modifications led to the emergence of epitranscriptome sequencing technologies profiling ribonucleotide modifications on a genomic scale ^21,22^. Here, we present a novel orthogonal chemistry-based sequencing strategy for the identification of 4-thiouridine (s^4^U), which is widely used as synthetic nucleotide-analog for in vivo, ex vivo and in vitro RNA labeling and represents a natural base modification in eubacterial and archaeal tRNA^12-16,42^. We show that chemical derivatization of s^4^U by thiol-linked alkylation induces reverse-transcriptase-dependent T>C conversions in sequencing data at single-nucleotide resolution and with a recovery rate of ≥90% (Fig.1 and Supplementary Fig. 9). At the same time, error rates in high-throughput sequencing approaches, e.g. using the Illumina platform, are low (<10^-3^, Fig. 2c), resulting in an overall signal-to-noise ratio of >900:1, which represents a significant improvement over previously described chemistries that are typically employed for chemo-selective s^4^U-enrichment by reversible biotinylation^15^.

By combining thiol-linked alkylation with s^4^U-metabolic RNA labeling in mouse embryonic stem cells, we show that SLAM-seq provides rapid access to intracellular kinetics of RNA biogenesis and decay in a global and transcript-specific manner (Fig.3,4 and 5). To this end we combined SLAM-seq with mRNA 3′ end sequencing (Quant-seq), a rapid workflow for the profiling of polyadenylated RNA polymerase II transcripts^24^. This approach provides several advantages with important practical and conceptual implications: (1) The specific sampling of poly-adenylated RNA species enables to assign kinetics to functional, fully processed RNA polymerase II transcripts (Fig.2 and Supplementary Fig.6). (2) In combination with Quant-seq, SLAM-seq provides access to mRNA 3′ isoform-specific expression-dynamics depending on polyA-site usage that may vary in different cell types (Supplementary Fig. 7). (3) Because mRNA 3′ end sequencing produces one fragment per transcript, downstream data analysis is facilitated by eliminating the requirement to normalize for transcript length (Supplementary Fig.6). (4) Quant-seq produces highly reproducible results from as little as 100 pg total RNA without requirement for rRNA depletion, hence provides access to cellular systems for which starting material is limiting^24^. (5) High sequencing coverage across inherently U-rich 3′ UTRs facilitates the robust quantification of T>C conversions. Note, that Quant-seq restricts gene expression analysis to RNA polymerase II transcripts and fails to differentiate transcript variants such as splice-isoforms. In future, alternative sequencing methods may augment the applicability of SLAM-seq, because s^4^U-identification by sequencing is in principle compatible with any RNA library preparation method that involves a reverse transcription step.

Studying intracellular RNA kinetics by s^4^U-metabolic RNA labeling requires general and method-specific considerations to be taken into account: While s^4^U is absent from metazoan cellular RNA and biologically inert unless exposed to UV-light, s^4^U-incorporation was previously linked to rRNA processing defects in human cancer cells ^43^. Because s^4^U-uptake can vary between cell types, careful assessment of cell-type-specific toxicity is imperative to meet s^4^U-labeling conditions that do not affect gene expression or cell viability (Supplementary Fig.5)^13,44^. In mES cells non-toxic concentrations of 100 μM s^4^U result in a median s^4^U-incorporation of 2.29% across 8408 transcripts upon long-term metabolic labeling (i.e. 24h), corresponding to one s^4^U incorporation in every 43 uridines at steady-state labeling conditions (Fig. 2c and Supplementary Fig. 8c). Considering the U-content of mRNA 3′ UTRs (∼31% in mES cells), SLAM-seq recovers each s^4^U-labeled transcript at a probability of up to 35% or 70% in single-read 50 or 100 sequencing reactions, respectively, which enables labeled-transcript identification even in short s^4^U pulse labeling conditions (Fig.3). Furthermore, the ability to differentiate labeled from unlabeled transcripts in the context of total RNA provides rapid access to transcript-specific labeling stoichiometry, hence circumvents the need for spike-in strategies to determine biogenesis and turnover rates^16,20^. Note, that the ability to determine de novo synthesized transcripts will depend on (1) the s^4^U uptake kinetics of the chosen cellular system, (2) the overall transcriptional activity of the cell type and (3) the library sequencing depth. Hence, these parameters need to be taken into account when designing a SLAM-seq experiment, particularly when employing short s^4^U pulse labeling, where sequencing depth demands adjustments to the given cellular parameters. Furthermore, s^4^U-tagging approaches (i.e. 4sU-seq or TT-seq) may provide some advantage over SLAM-seq when analyzing transient RNA species that escape detection by standard RNA sequencing approaches^16^.

Finally, we compared SLAM-seq extracted rates to existing published estimates. We compared transcriptional output measurements generated by SLAM-seq to GRO-seq data and found a significant correlation for RNA polymerase II transcripts in mES cells (Supplementary Fig. S11)^29^. Median mRNA half-life in mES cells (4.3 h; Figure 4d) falls within the range of previously determined mRNA stabilities in mammalian cells, ranging in median half-life between 3.4h and 10h, depending on cell type and experimental technique (Tani et al., 2012). Furthermore, mRNA stabilities derived from SLAM-seq measurements exhibit an overall good correlation with half-life determined by global transcriptional inhibition (Supplementary Fig.14), and recapitulate previously proposed relationships between mRNA stability and its physiological function (Fig. 4d and Supplementary Fig.12)^1^.

In summary, SLAM-seq enables cost-effective insights into intracellular RNA kinetics at unprecedented accessibility, efficiency and scalability, hence represents a powerful tool for the dissection of fundamental biological mechanisms that control metazoan gene expression.

## Online Methods

### Carboxyamidomethylation of s^4^U

If not indicated otherwise, carboxyamidomethylation was performed under standard conditions (50% DMSO, 10 mM iodoacetamide, 50 mM sodiumphosphate buffer pH8, for 15 min at 50°C) using either 1 mM 4-thiouracil (Sigma), 800 μM 4-thiouridine (Sigma), or 5 – 50 μg total RNA prepared from s^4^U metabolic labeling experiments. The reaction was quenched by addition of excess DTT.

### Adsorption measurements

1mM 4-thiouracil was incubated under optimal reaction conditions (10mM iodoacetamide, 50% DMSO, 50 mM sodiumphosphate buffer pH8, for 15 min at 50°C) if not indicated otherwise. Reaction was quenched by the addition of 100 mM DTT and adsorption spectra were measured on a Nanodrop 2000 instrument (Thermo Fisher Scientific), followed by baseline subtraction of adsorption at 400 nm.

### Mass Spectrometry

40 nmol 4-thiouridine were reacted in the absence or presence of 0.05, 0.25, 0.5 or 5 μmol iodoacetamide under standard reaction conditions (50 mM sodiumphosphate buffer, pH 8; 50 % DMSO) at 50°C for 15 minutes. The reaction was stopped with 1% acetic acid. Acidified samples were separated on a Ultimate U300 BioRSLC HPLC system (Dionex; Thermo Fisher Scientific), employing a Kinetex F5 Pentafluorophenyl column (150 mm x 2.1 mm; 2.6 μm, 100 Å; Phenomenex) with a flow rate of 100 μl/min. Nucleosides were on-line analyzed using a TSQ Quantiva mass spectrometer (Thermo Fisher Scientific) after electrospray ionization with the following SRMs: 4-Thiouridine m/z 260 → 129, and alkylated 4-Thiouridine m/z 318 → 186. Data were interpreted using the Trace Finder software suite (Thermo Fisher Scientific) and manually validated.

To determine s^4^U incorporation events in polyadenylated or total RNA by Mass Spectrometry, total RNA was either subjected to oligo(dT) enrichment using Dynabeads® Oligo(dT)25 (Ambion) following manufacturer’s instructions to purify polyadenylated RNA or directly enzymatically degraded to monomeric ribonucleosides as described previously prior to Mass Spectrometry analysis^45^.

### Primer extension assays

Primer extension assays were essentially performed as described previously^46^. Briefly, template RNA oligonucleotides (5L-*let-7*-3L or 5L-*let-7*-s^4^Up9-3L; Dharmacon; see Supplementary Table 3 for sequences) were deprotected according to the instructions of the manufacturer and purified by denaturing polyacrylamide gel-elution. 100 μM purified RNA oligonucleotides were treated with 10 mM iodoacetamide (+IAA) or EtOH (-IAA) in standard reaction conditions (50 % DMSO, 50 mM sodiumphosphate buffer, pH8) for 15 min at 50°C. The reaction was stopped by addition of 20 mM DTT, followed by ethanol precipitation. RT primer (see Supplementary Table 3 for sequence) was 5′ radiolabeled using γ-^32^P-ATP (Perkin-Elmer) and T4-polynucleotide kinase (NEB), followed by denaturing polyacrylamide gel-purification. 640 nM γ-^32^P-RT primer was annealed to 400 nM 5L-*let-7*-3L or 5L-*let-7*-s^4^Up9-3L in 2 x annealing buffer (500 mM KCl, 50 mM Tris pH 8.3) in a PCR machine (3 min 95°C, 30 sec 85°C Ramp 0.5°C/s, 5 min 25°C Ramp 0.1°C/s). Reverse transcription was performed using Superscript II (Invitrogen), Superscript III (Invitrogen), or Quant-seq RT (Lexogen) as recommended by the manufacturer. For dideoxynucleotide reactions, a final concentration of 500 μM ddNTP (as indicated) was added to RT reactions. Upon completion, RT reactions were resuspended in formamide loading buffer (Gel loading buffer II, Thermo Fisher Scientific) and subjected to 12.5% denaturing polyacrylamide gel electrophoresis. Gels were dried, exposed to storage phosphor screen (PerkinElmer), imaged on a Typhoon TRIO variable mode imager (Amersham Biosciences), and quantified using ImageQuant TL v7.0 (GE Healthcare). For analysis of RT drop-off, signal-intensities at p9 were normalized to preceding drop-off signal intensities (bg, Supplementary Fig.3b) for individual reactions. Values reporting the change in drop off signal (+IAA/-IAA) for s^4^U-containing and non-containing RNA oligonucleotides were compared for the indicated reverse transcriptases.

## HPLC analysis of s^4^U-labeled RNA

Analysis of s^4^U-incorporation into total RNA following metabolic labeling was performed as previously described^45^.

### Cell viability assay

5000 mES cells were seeded per 96 well the day before the experiment. After onset of the experiment, media containing the indicated concentration of s^4^U was replaced every three hours for a total of 12 h or 24 h. Cell viability was assessed by CellTiter-Glo® Luminescent Cell Viability Assay (Promega) according to the instructions of the manufacturer. Luminescent signal was measured on Synergy (BioTek) using Gen5 Software (v2.09.1).

### Cell culture

Mouse embryonic stem (mES) cells (clone AN3-12), derived from C57BL/6×129 F1 females, were obtained from IMBA Haplobank (U. Elling et al., accepted for publication in Nature) and cultured in 15 % FBS (Gibco), 1× Penicillin-Streptomycin solution (100 U/ml Penicillin, 0.1 mg/ml Streptomycin, Sigma), 2 mM L-Glutamine (Sigma), 1× MEM Non-essential amino acid solution (Sigma), 1 mM sodium pyruvate (Sigma), 50 μM 2-Mercaptoethanol (Gibco) and 20 ng/ml LIF (in-house produced). Cells were maintained at 37°C with 5% CO_2_ and passaged every second day. Cell doubling time of AN3-12 mES in presence of s^4^U cells as determined by cell counting was 14.7h. Prior to metabolic labeling experiments, mES cells were stained with Hoechst33342 and FACS-sorted to obtain a pure diploid population^47^.

### SLAM-seq in mES cells

mES cells were seeded the day before the experiment at a density of 10^5^ cells/ml. s^4^U-metabolic labeling in mES cells was performed by incubating mES cells in standard medium but adding s^4^U (Sigma) to a final concentration of 100 μM and media exchange every 3 hours for the duration of the pulse. For the uridine chase experiment, cells were washed twice with 1x PBS and incubated with standard medium supplemented with 10 mM uridine (Sigma). At respective time points, cells were harvested followed by total RNA extraction using TRIzol (Ambion) following the manufacturer’s instructions but including 0.1mM DTT (final conc.) during isopropanol precipitation. RNA was resuspended in 1 mM DTT. For a typical SLAM-seq experiment, 5 μg total RNA were treated with 10 mM iodoacetamide under optimal reaction conditions and subsequently ethanol precipitated and subjected to Quant-seq 3′ end mRNA library preparation.

### RNA library preparation

Standard RNA seq libraries were prepared using NEBNext® Ultra(tm) Directional RNA Library Prep Kit for Illumina® (NEB) following the instructions of the manufacturer. Cap-seq libraries were prepared as previously described^48^. mRNA 3′ end sequencing was performed using the Quant-seq mRNA 3′ end library preparation kit (Lexogen) according to the instructions of the manufacturer. Small RNA libraries were generated as described before^49^, but adding total RNA from Arabidopsis thaliana unopened floral buds as spike-in before initial size-selection. Sequencing was performed on Illumina HiSeq 2500. Libraries were sequenced in SR50 mode except for transcriptional output measurements (Fig.3), which were sequenced in SR100 mode.

### Transcriptional inhibition by Actinomycin D

3 × 10^5^ AN3-12 mES cells were seeded per 35 mm plate and grown over night. To block transcription, actinomycin D (Sigma) was added to the medium at the concentration of 5 μg/ml. Cells were harvested at 0, 0.25h, 0.5h, 1h, 3h and 10 h after addition of actinomycin D by directly lysing them in TRIzol® (Ambion). RNA was extracted following the manufacturer instructions and libraries were prepared using Quant-seq mRNA 3′ end library preparation kit (Lexogen) according to the instructions of the manufacturer.

### CRISPR/Cas9 genome engineering

gRNAs were designed using WTSI Genome Editing^50^. gRNA oligonucleotides (see Supplementary Table 3) were cloned into pLenti-CRISPR-v2-GFP vector as described^51^, but modified by replacing the puromycin resistance cassette with GFP. Prior to gRNA transfection targeting Xpo5 or Mettl3, wildtype An3-12 mES cells were FACS sorted for haploid cells as described previously^47^. 3 × 10^5^ cells were seeded per 6 well and transfected the next day with 3 μg pLenti-CRISPR-v2-GFP using Lipofectamine 2000 as recommended by the manufacturer. 48h after transfection, GFP positive cells were sorted by fluorescence-activated cell sorting (FACS) and 1500 cells were subsequently seeded per 15 cm plate. Single colonies were picked after 10 days. DNA isolation, PCR amplification (for oligonucleotide sequences see Supplementary Table 3) of the targeted locus and Sanger sequencing was performed to genotype the clonal cell lines. Protein depletion was confirmed by Western blot analysis.

### Western Blotting

Protein lysates were separated on 10% SDS PAGE and transferred to PVDF membrane (BioRad). Antibodies were used at a dilution of 1:500 for anti-Exportin-5 (H-300, sc-66885, rabbit), 1:3,000 for anti-Mettl3 (15073-1-AP, Proteintech, rabbit), 1:5,000 for anti-Mettl14 (HPA038002, Sigma, rabbit) and 1:10,000 for anti-Actin (A2066, Sigma, rabbit) and detected by secondary HRP-antibody-conjugates G21040 (Invitrogen; dilution 1:10,000). Primary antibodies were incubated at room temperature for three hours and secondary antibodies were incubated at room temperature for two hours. Images were acquired on a ChemiDoc MP Imaging System (BioRad) using ImageLab v5.1.1 (BioRad) or by Amersham Hyperfilm ECL (GE Healthcare).

### Northern Blotting

Northern hybridization experiments were performed as described previously^52^. For Northern probes see Supplementary Table 3.

### Bioinformatics and Data analysis

Gel images were quantified using ImageQuant v7.0a (GE Healthcare). Curve fitting was performed according to the integrated rate law for a first-order reaction in Prism v7.0 (GraphPad) or R (v2.15.3) using the minpack.lm package. Statistical analyses were performed in Prism v7.0a (GraphPad), Excel v15.22 (Microsoft) or R (v2.15.3 and v3.3).

For sequencing analysis of synthetic RNA samples (Fig.1i and Supplementary Fig.4) barcoded libraries were demultiplexed using Picard Tools BamIndexDecoder v1.13 allowing 0 mismatches in the barcode. Resulting files were converted to fastq using picard-tools SamToFastq v1.82. Cutadapt v1.7.1 was used to trim adapters (allowing for default 10% mismatch in adapter sequence) and filter for sequences of 21nt length. Resulting sequences were aligned to aligned to mature dme-let-7 sequence (TGAGGTAGTAGGTTGTATAGT) using bowtie v0.12.9 allowing for 3 mismatches and converted to bam using samtools v0.1.18. “N” containing sequences were filtered from alignment. Remaining alignments were converted to pileup format. Finally, fraction of each conversion per position were extracted from pileup. Output table was analyzed and plotted in Excel v15.22 (Microsoft) and Prism v7.0a (GraphPad).

For standard RNA sequencing data analysis, barcoded libraries were demultiplexed using Picard Tools BamIndexDecoder v1.13 allowing 1 mismatch in the barcode. Adapters were clipped using cutadapt v1.5 and reads were size-filter for ≥ 15 nucleotides. Reads were aligned to mouse genome mm10 using STAR aligner v2.5.2b^53^. Alignments were filtered for alignment scores ≥ 0.3 and alignment identity ≥ 0.3 was normalized to read length. Only alignments with ≥ 30 matches were reported and chimeric alignments with an overlap ≥ 15 bp were allowed. 2-pass mapping was used. Introns < 200 kb were filtered and alignments containing non-canonical junctions were filtered. Alignment with a mismatch to mapped bases ratio ≥ 0.1 or with a max. number of 10 mismatches were excluded. The max number of gaps allowed for junctions by 1,2,3,N reads was set to 10 kb, 20kb, 30kb and 50 kb, respectively. The minimum overhang length for splice junctions on both sides for (1) non-canonical motifs, (2) GT/AG and CT/AC motif, (3) GC/AG and CT/GC motif, (4) AT/AC and GT/AT motif was set to 20, 12, 12, 12, respectively. “Spurious” junction filtering was used and the maximum number of multiple alignments allowed for a read was set to 1. Exonic reads (Gencode) were quantified using FeatureCounts^54^.

For Cap analysis gene expression (Cap-Seq), barcoded libraries were demultiplexed using Picard Tools BamIndexDecoder v1.13 allowing 1 mismatch in the barcode. The first 4nt of the reads were trimmed using seqtk. Reads were screened for ribosomal RNA by aligning with BWA (v0.6.1)^55^ against known rRNA sequences (RefSeq). The rRNA subtracted reads were aligned with TopHat (v1.4.1)^56^ against the Mus musculus genome (mm10). Maximum multihits was set to 1, segment-length to 18 and segment-mismatch to 1. Additionally, a gene model was provided as GTF (Gencode VM4).

For analysis of mRNA 3′ end sequencing (Quant-seq) datasets, reads were demultiplexed using Picard Tools BamIndexDecoder v1.13 allowing 1 mismatch in the barcode. Quant-seq data was processed using Digital Unmasking of Nucleotide conversion-containing k-mers (DUNK), SLAM-DUNK v0.2.4, a T>C aware alignment software package based on NextGenMap^57^ developed to recover T>C conversions from SLAM-seq data sets (Neumann T., *et al.*, in preparation). Briefly, adapter-clipped reads were trimmed 12 bp from the 5′ end (-5 12) and poly(A) stretches (>4 subsequent As at the 3′ end) were removed. Trimmed reads were aligned to the full reference genome (mm10) using local alignment scoring and up to 100 alignments were reported for multimapping reads (-n 100). In the filtering step, alignments with a minimum identity of 95% and a minimum of 50% of the read bases mapped were retained. Among multimappers, reads mapping to no or ambiguously to > 1 annotated UTR sequence (bed files provided in GEO datasets) were discarded (-fb). If a multimapping read mapped >1 time to the same annotated UTR sequence, one alignment was randomly picked. SNPs exceeding a coverage cutoff of 10x and a variant fraction cutoff of 0.8 were called using VarScan2.4.1 using default parameters^58^. Non-SNP overlapping T>C conversions with a base quality of Phred score >26 were identified. T>C containing reads and total reads aligning within the custom defined counting windows (bed files provided in GEO datasets) were reported. T>C conversion rate was determined for each position along the custom defined counting windows by normalizing to genomic T content and coverage of each position and averaged per UTR.

For extended mRNA 3′ end annotation, we assembled a pipeline to annotate 3′ ends of mRNA transcripts using Quant-seq datasets (https://github.com/AmeresLab/UTRannotation). Quant-seq data was pre-processed as described above. To determine exact priming sites, reads with continuous 3′ terminal A stretches (> 4) and a length of at least 23 nts long were retained. Polymeric A-stretches were trimmed from the 3′ ends of reads and mapped to mm10 using SLAM-DUNK’s map and filter module as described above but using global alignment scoring. Priming sites were identified based on mapping of >= 10 reads to genomic positions and consecutive positions were merged. Genomic A content of >= 0.36 and >=0.24 was used to identify internal priming events (for polyA site-containing and no-polyA site-containing priming sites respectively, see Supplementary Fig.7 for PAS sequences). Priming sites overlapping with RefSeq and ENSEMBL 3′ UTR annotations were considered for further analysis (UTRends). RNA-seq signal, mapped as described above, was used to identify intergenic ends. RNA-seq coverage was calculated using bedtools multicov in 200nt bins separated by 20nts starting from the last 200nts of gene annotations. Bins were extended until RNA-seq coverage dropped below 10% compared to the first bin or until the bin overlapped another gene annotation. Priming sites overlapping identified counting bins were retained (intergenicEnds). For each gene, all identified 3′ ends were ranked by underlying counts and ends that did not exceed 10% of the total signal were removed. RefSeq-annotated mRNA 3′ ends were then included and 250nt counting windows were created upstream of 3′ ends. Overlapping counting windows were merged. Beyond protein coding mRNAs, counting windows were added for the following classes of non-coding RNAs: antisense, bidrectional_promoter_lncRNA, lincRNA, macro_lncRNA, processed_transcript, sense_intronic, sense_overlapping and primary miRNAs. To annotate 3′ UTR start positions for de-novo annotated 3′ ends, each 3′ end was assigned to the most proximal 3′ UTR start annotation (RefSeq).

For comparison of Quant-seq and RNA-seq, we employed RefSeq transcripts of mm10 from UCSC’s table browser (downloaded 2017-02-14) consisting of 35,805 transcripts which we mapped to 24,440 Entrez genes. All transcripts for a given gene were merged using bedtools^59^. Stranded coverage tracks for Quant-seq and RNA-seq samples were created using deeptools’ *bamCoverage* command^60^, using a binSize of 1 and normalizing to RPKM. Next, the density matrix was calculated separately for + and – strand genes, with static windows 500 bp in both directions at TSS and TTS and dynamic binning for the remaining gene body. Stranded signal from the density matrix was plotted in composite plots.

For transcriptional output analysis, the number of normalized reads (in cpm; “Steady-state Expression”) and the number of normalized reads containing ≥1 T>C conversion (in cpm; “Transcriptional Output”) were obtained for every gene after aligning SLAM-seq data with SLAM-DUNK to the mouse genome mm10. Background T>C reads (T>C reads observed without s^4^U labeling) were subtracted from the T>C reads in the 45min time-point and an expression threshold of >5 cpm for the mean of “Steady-state Expression” was set. Genes were classified as proximal to “no”, “OSN” or “strong/super” enhancer according to Whyte et al.^27^.

GRO-seq data from mES cells was downloaded from GEO (GSE27037)^29^. Reads were mapped to mm10 using bowtie allowing for uniquely mapping reads with at most 2 mismatches. Unmapped reads were reiteratively trimmed by one nucleotide and remapped until reaching a minimum length of 20 nucleotides. GRO-seq signal was assessed using featureCounts^54^ for the full length gene omitting the first kilobase. Transcriptional output as determined by SLAM-seq was then compared to GRO-seq for all genes that are expressed above 5cpm in Quant-seq datasets and detected in GRO-seq datasets.

To calculate RNA half-lives, T>C conversions were background-subtracted (no s^4^U treatment) and normalized to chase-onset. Curve fitting was performed according to the integrated rate law for a first-order reaction in R (v2.15.3) using the minpack. lm package. RNA half-lives > 24h were set to 24h. If not stated otherwise an R^2^ cutoff of > 0.6 was applied. To calculate RNA half-lives normalized to cell cycle length, T>C conversions were multiplied by 2^(timepoint/14.7h)^.

To calculate RNA stabilities measured by polymerase II inhibition (ActD treatment), reads from the Actinomycin D-treated samples were aligned to mm10 using SLAM-DUNK. Transcripts were extracted that were expressed > 5cpm in the SLAM-seq experiment. To correct for the relative increase in stable transcripts following global transcriptional inhibition, data was normalized to the 50 most stable transcripts. Half-lives were calculated by fitting data to a single-exponential decay model as described above.

GO terms-enrichment analysis was performed using PANTHER database with a custom reference set consisting of genes expressed > 5cpm in mES cells (n=8533)^61^. For gene-set enrichment analysis, gene-association with GO terms “Regulation of Transcription” (GO:0006357), “Cell cycle” (GO:0007049), “Translation” (GO:0006412) and “Extracellular Matrix” (GO:0031012) were derived from AmiGO^62^.Transcripts were pre-ranked based on the difference half-life to the mean half-life after log_2_-transformation. GSEAPreranked was performed using GSEA v.2.2.4^63,64^.

MicroRNA targets were predicted using Targetscan v7^65^. Briefly, we provided a 60-way multiple genome alignment against mm10 and our custom 3′-end annotation to create a tailored database of conserved miRNA targets. The output was then intersected with our data, filtered, and grouped according by site type. To determine site conservation, cutoffs for branch length score were set to ≥ 1.6 (“7mer-1a”), ≥ 1.3 (“7mer-m8”) and ≥ 0.8 (“8mer”).

Relative RNA stabilities were determined by performing SLAM-seq after 3h and 12h s^4^U pulse labelling in wildtype or knock-out cell lines. The background subtracted T>C conversion rates at 3h were normalized to 12h and relative stabilities for control (treated with non-targeting gRNA^51^) and knockout cells were assessed from the following equation: ln(2) / ln(1-(T>C conversion [3h] / T>C conversion [12h]))/3.

N6-methyladenosine-targets were extracted from Batista et al., 2014^37^ and batch coordinate conversion (liftOver) from mm9 to mm10 (UCSC) was performed, or from Ke et al., 2017^41^. Tags in 3′ UTRs were refined by overlapping the genomic coordinates with the custom mES cell annotation.

## Accession code

Sequencing data associated with this manuscript is available at GEO under the accession number GSE99978.

## Acknowledgements

We thank J. Jude for generously providing a modified version of pLenti-CRISPR-v2-GFP, Gabriela Krssakova for HPLC analysis, the IMP/IMBA Biooptics facility for FACS support, and all laboratory members for support and discussions. Mass spectrometry was performed at the VBCF Metabolomics unit (www.vbcf.ac.at), funded by the City of Vienna through the Vienna Business Agency. HTP sequencing was performed at the VBCF NGS Unit (www.vbcf.ac.at). This work was supported by the Austrian Academy of Sciences, the Austrian Science Fund FWF (Y-733-B22 START, W127-B09, and F4322), and the European Research Council (ERC-StG-338252 miRLIFE) to S.L.A

